# Anti-Bat Ultrasound Production in Moths is Globally and Phylogenetically Widespread

**DOI:** 10.1101/2021.09.20.460855

**Authors:** JR Barber, D Plotkin, JJ Rubin, NT Homziak, BC Leavell, P Houlihan, KA Miner, JW Breinholt, B Quirk-Royal, PS Padrón, M Nunez, AY Kawahara

## Abstract

Warning signals are well known in the visual system, but rare in other modalities. Some moths produce ultrasonic sounds to warn bats of noxious taste or to mimic unpalatable models. Here we report results from a long-term study across the globe, assaying moth response to playback of bat echolocation. We tested 252 genera, spanning most families of large-bodied moths, and outline anti-bat ultrasound production in 52 genera, with eight new subfamily origins described. Based on acoustic analysis of ultrasonic emissions and palatability experiments with bats, it seems that acoustic warning and mimicry are the *raison d’etre* for sound production in most moths. However, some moths use high-density ultrasound capable of jamming bat sonar. In fact, we find preliminary evidence of independent origins of sonar jamming in at least six subfamilies. Palatability data indicates that jamming and warning are not mutually exclusive strategies. To explore the possible organization of anti-bat warning sounds into acoustic mimicry rings, we intensively studied a community of moths in Ecuador and found five distinct acoustic clusters using machine learning algorithms. While these data represent an early understanding of acoustic aposematism and mimicry across this megadiverse insect order, it is likely that ultrasonically-signaling moths comprise one of the largest mimicry complexes on earth.

## Introduction

Across systems, unpalatable prey declare their location and identity to predators (1). Gaudy poison frogs and red newts alert attackers of toxins sequestered in their skin glands (2, 3), brightly banded coral snakes warn birds of their venomous bite (4), and patterned milkweed bugs and monarch butterflies proclaim their unpalatable hemolymph (5). While aposematism (conspicuous signaling to advertise noxiousness (6)) has been most rigorously studied in the visual system, warning displays have also been described in the olfactory (7) and auditory systems (8). Until now, acoustic aposematism has appeared as either an accessory in a multi-sensory warning suite (9), or a highly specialized and unique antipredator trait (8, 10). Here, we describe one of the world’s largest and most widespread aposematic complexes: ultrasonic clicking by chemically-defended nocturnal moths and their purported mimics.

Moths fly in a dim, acoustic world. Over millions of years they have repeatedly evolved ears (11), organs that likely originated for general auditory surveillance of the environment (12), and that were secondarily co-opted to detect the sonar cries of bats. Hearing organs are found in many regions of the lepidopteran body and occur in a significant majority of species in the order (including ∼85% of species in the megadiverse Macroheterocera) (13–15). These advance warning sensors allow moths to hear echolocating bats and either motorically evade attack by steering away or performing acrobatic loops, spirals and dives (16), or respond to bats with a countervailing signal of their own. Ultrasonic clicking by moths, in response to bat sonar, has been documented in tiger moths (17), hawkmoths ((18, 19), and one geometrid moth (20). These sounds can function non-mutually-exclusively to jam bat sonar (18, 21, 22), signal noxiousness (or mimic noxious acoustic models) (8, 23), and startle bat predators (24).

We hypothesized that, given the efficacy of anti-bat ultrasound production by moths in the hawkmoth and tiger moth lineages, sound emission was perhaps common and widespread across the entire order of more than 160,000 described lepidopteran species. Here, we report a long-term dataset from research across the globe, assaying moth response to playback of bat attack. We tested 252 genera, spanning most families of relatively large-bodied moths (i.e., exceeding 1 cm in length and/or wingspan), and describe anti-bat sound production in 52 genera (21%). For most of these genera, this is novel behavior never before described. This number is a clear underestimate of acoustic aposematism, mimicry, and sonar jamming across this megadiverse insect order (1 in 10 described animals on Earth is a lepidopteran (25)).

## Results and Discussion

To uncover the prevalence of ultrasonic response to echolocating bat attack, we trapped moths with UV lights and broadcast pre-recorded bat sonar attack sequences to moths in tethered flight, across the world’s tropics from Asia and Africa (Malaysian Borneo and Mozambique) to South America (Ecuador, and French Guiana). Using an ultrasonic speaker, we played representative calls from species of both frequency-modulated (FM; characterized by short-duration, frequency-sweeping pulses) and constant-frequency (CF; characterized by tonal, long-duration pulses (26)) bats (see Fig. S1). We recorded moth responses to playback of sonar attack and found that 52 of 252 tested genera respond acoustically to both types of bat sonar (Fig. 1, Dataset S1, Supp. Archive 10) – discoveries that now add nine subfamilies to those known to employ this defense (19, 27, 28). While anti-bat ultrasound has been described and well-studied in arctiines (tiger moths) (28–30) and sphingids (hawkmoths) (18, 19, 31), here we report that this striking anti-predator behavior is widespread across the tapestry of lepidopteran diversity (Fig 2). In fact, if we extrapolate from our sample, ∼20% of the estimated 100,000 species of Macroheterocera (12) produce ultrasound in response to bat sonar.

**Figure 1.**
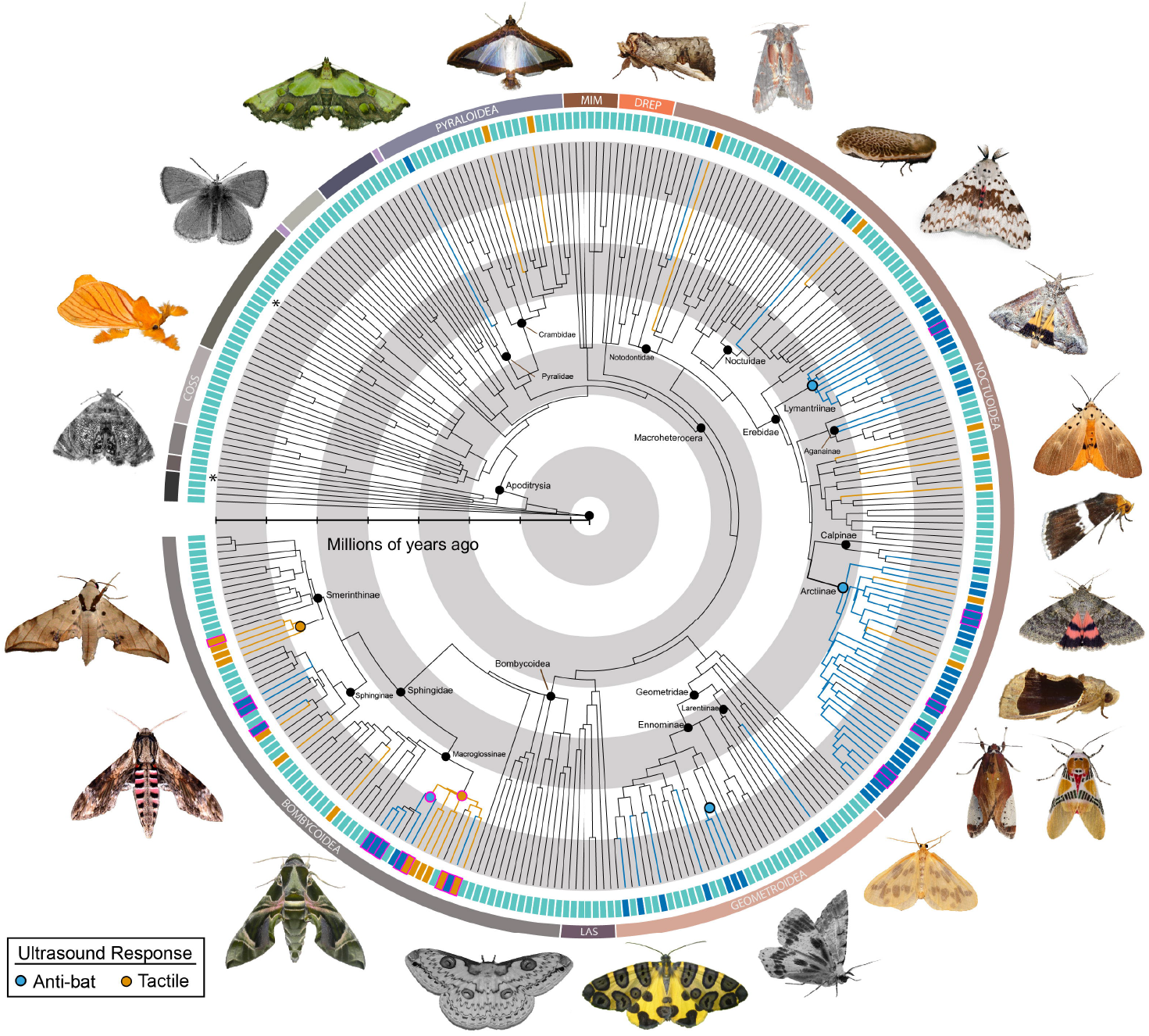
A molecular phylogeny of Lepidoptera indicating anti-predator ultrasound production across the order. Bars and nodes with magenta outlines represent taxa associated with sufficiently large duty cycle values (>18%) for sonar jamming. Asterisks indicate taxa known to produce ultrasound, but not in response to either tactile stimuli nor bat ultrasound. Grayscale images indicate taxa that do not produce ultrasound. This phylogeny is meant to illustrate the diversity of ultrasound production and offer broad strokes on the origins of anti-predator sounds at the family and subfamily level, not as a test of evolutionary relationships. Photographs are distributed under Creative Commons Attribution NonCommercial Licenses (see Fig. S2, Dataset S3 for full accreditations).

**Figure 2.**
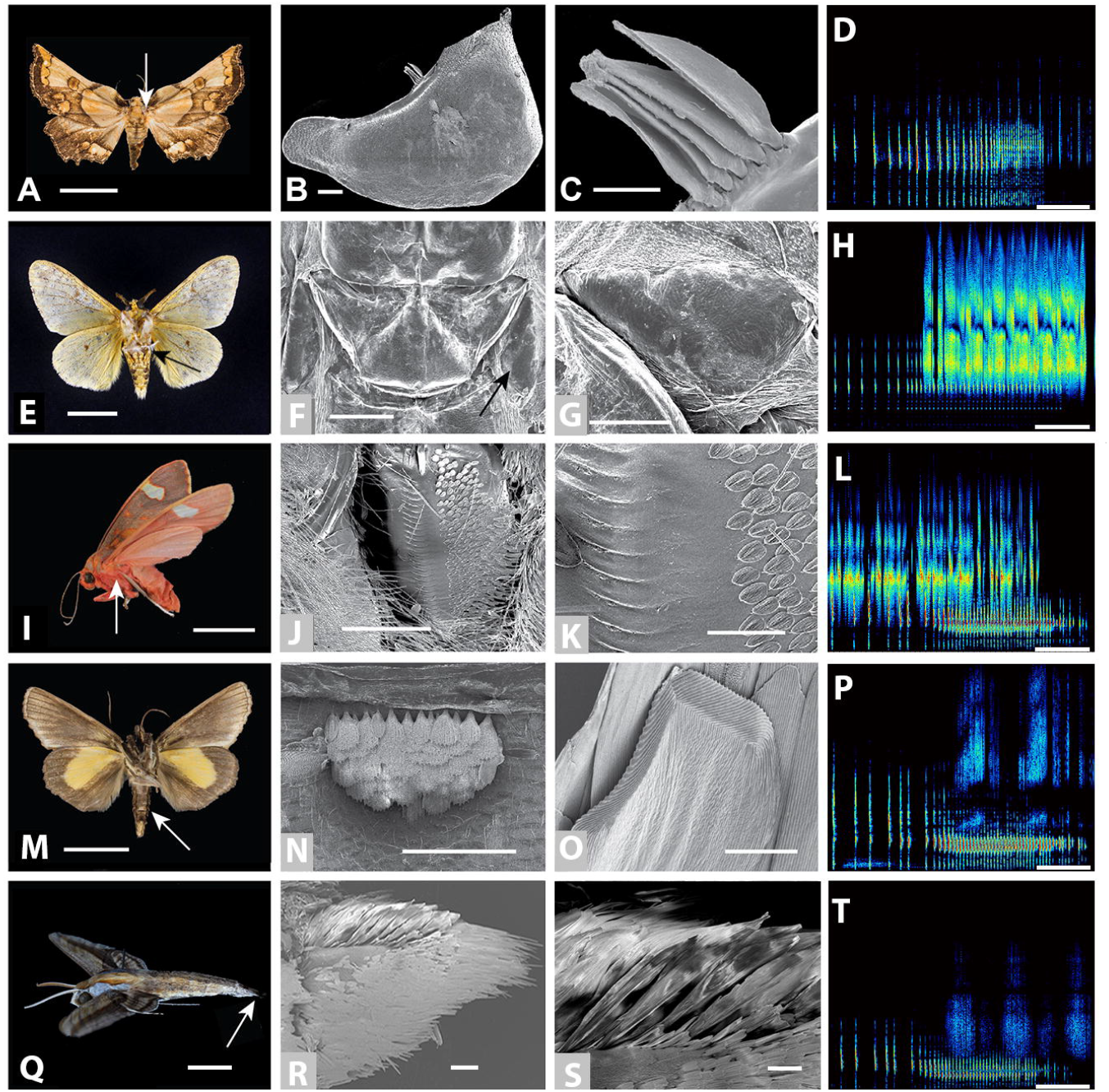
Anti-bat ultrasound-producing structures. A-D. *Mittonia hampsoni* (Pyralidae: Pyralinae) produces ultrasonic clicks in flight via modified scales on the tegula; A. Scale bar = 1.0 cm; B. Tegula, 0.2 mm; C. Tegular scales, 50 μm; D. Response to bat sonar playback (*Mittonia hampsoni*), 100 ms). E-H. *Lymantria sp*. (Erebidae: Lymantriinae) generates ultrasound with paired tymbals recessed in abdominal pockets; E. Scale bar = 1.0 cm; F. Arrow indicates one of the tymbal pair, 1.0 mm; G. Close up of one tymbal, 0.5 mm; H. Response to bat sonar playback (*Lymantria sp*.), 100 ms. I-L. *Melese sp*. (Erebidae: Arctiinae) emits ultrasound with paired thoracic tymbals; I. Scale bars = 1.0 cm; J. Tymbal 0.5 mm; K. Close-up of microstriations on tymbal surface, 0.1 mm; L. Response to bat sonar playback (*Melese peruviana*), 100 ms. M-P. *Gonodonta sicheas* (Erebidae: Calpinae) produces ultrasound by stridulating modified abdominal scales; M. Scale bar = 1.0 cm; N. Patch of stridulatory scales, 0.5 mm; O. Stridulatory scale, Scale bar = 50 μm; P. Response to bat sonar playback (*Gonodonta bidens*), 100 ms. Q-T. *Xylophanes falco* (Sphingidae: Macroglossinae) produces ultrasound by stridulating modified genital valves; Q. Scale bar = 1 cm; R. Patch of stridulatory scales on genital valve, 0.5 mm; S. Stridulatory scales, 0.2 mm; T. Response to bat sonar playback (*Xylophanes amadis*), 100 ms.

In addition to playback of bat attack, we also queried moths for ultrasonic response to handling. We simulated a physical predatory attack by grasping the thorax, abdomen, and head. Nearly all moth species that broadcast anti-bat sounds upon hearing sonar also produced ultrasonic disturbance sounds when handled. Three subfamilies from three different families (Erebidae: Erebinae, Crambidae: Spilomelinae, Sphingidae: Smerinthinae; see Dataset S2) produced ultrasound only in response to tactile stimulation. Producing ultrasound to touch may be a generalized anti-predator response intended to startle attackers (32). Moreover, responding to bats during handling may still provide time for bats to recognize the warning signal and drop these moths unharmed (*sensu* (27)), as bats often first contact their prey with an outstretched wing, directing the insect to their tail membrane, and then subsequently to their mouth (33). Indeed, in a study that pit northern long-eared bats (*Myotis septentrionalis*) against aposematically clicking dogbane tiger moths (*Cycnia tenera*), 75% of signaling moths that were captured were subsequently dropped unscathed (34). The critical experiments pitting bats against moths that produce ultrasound to physical contact only have yet to be performed.

Our data indicate that ultrasound production has arisen repeatedly in novel and convergent forms. To determine the mechanism of ultrasonic clicking in each newly discovered sound producer, we recorded synchronized audio and macro medium-speed video (∼100 fps) footage of moths producing ultrasound (see Movies S1–S2). We found several different mechanisms across and within lineages, and a great deal of morphological convergence (Fig. 2). The sound-producing mechanisms we uncovered can be grouped into three broad categories: 1) abdominal stridulation, where modified scales on adjoining areas of the moth form a file-scraper device (e.g., Sphingidae: Macroglossinae, Sphingidae: Sphinginae, Erebidae: Calpinae); 2) percussive wing beating, where sound is produced on each wing stroke by moving the tegula into a striking position between the beating wings (e.g., Pyralidae: Pyralinae); and 3) tymbals, where thin, striated cuticular plates buckle under muscular force and passively release often making a series of clicks during each action due to striations on the tymbal’s surface (e.g., Erebidae: Lymantriinae, Erebidae: Aganainae, Erebidae: Arctiinae).

Previous work has shown that tiger moths (Erebidae: Arctiinae) and hawkmoths (Sphingidae) use tymbals and stridulation, respectively, to produce ultrasound in response to echolocating bat attack (18, 21, 27). Here we describe three new mechanisms of ultrasound production (Fig. 2): one stridulation-based, one tegula-based, and one tymbal-based. Calpines (a subfamily within Erebidae, here represented by the genus *Gonodonta*) stridulate using modified ventral abdominal scales (see Fig. 2M-P, Movie S1) that produce remarkably similar sounds to sphingids, which stridulate with modified scales on the genital valves (18, 19); Fig. 2Q-T). We found the percussive wing beating strategy in only one pyralid moth, *Mittonia hampsoni*, that facultatively beats its wings against its tegula (a structure that plays a role in protecting the base of the forewing; Fig. 2A-D) in flight, which we confirmed via ablation experiments. Lymantriines (Erebidae) use paired abdominal tymbals hidden within pockets that form horn-like structures when opened (see Fig. 2E-H, Movie S2), beaming ultrasound backwards at attacking bats.

Aganaines (Erebidae) use paired metathoracic tymbals in the identical positions to arctiines, calling into question the tymbal as a uniting characteristic of arctiines (tiger moths) (35, 36). Previous work described a geometrid (Geometridae: Larentiinae) that uses prothoracic tymbals to generate ultrasonic warning sounds (37). Here we discovered that multiple genera in a different geometrid subfamily, Ennominae, also produce anti-bat emissions. We have been unable to find a prothoracic tymbal in this group, presenting the intriguing possibility that anti-bat sound production has originated independently at least twice in geometrids. Despite our efforts in the field and museum, there are several other moth subfamilies in which we have confirmed ultrasound production for which we do not know the underlying mechanism (Crambidae: Spilomelinae, Erebidae: Erebinae, Erebidae: Hypocalinae, Noctuidae: Hadeninae, Noctuidae: Noctuinae, Notodontidae: Notodontinae, Notodontidae: Nystaleinae). Clearly, the mechanisms driving the acoustic arms race between moths and bats are myriad and diverse.

We also discovered an interesting form of ultrasound production in the Dalceridae (genus *Acraga*). These non-eared animals constantly produce ultrasound while in flight similar to the behaviors previously described in other small-bodied non-Macroheterocera (38, 39). The mechanism of sound production in the *Acraga* genus remains unknown – the wing-based aeroelastic tymbals implicated in sound production in other non-Macroheterocera do not appear responsible. Considering that moths in the genus *Acraga* are unpalatable to bats (see supplement), it is tempting to assert that these sounds are involved in advertising noxious taste to echolocating bats. Until moths using this type of ultrasound production are pit against bats in appropriate experiments, the function of these sounds will remain unclear.

To better understand how the interactions between bats and sound-producing moths might play out across the night skies, we quantified moth acoustic emissions, using previously-described parameters to capture the temporal and spectral components (27). We found that animals that produce ultrasound to playback of bat attack emit frequencies centered around ∼65 kHz (± ∼40-110 kHz at 15 dB range; matching the frequency of best hearing in most bat species (40, 41)) and a substantial range of duty cycles (sound per unit time; see Supp. Archive S10). While it is possible that any duty cycle sound can startle naive bats, or warn of noxious taste (or mimic chemically-protected models), only high duty cycle sounds can jam bat sonar (8, 10, 18, 22, 42, 43). In fact, duty cycles of at least 18% (this value is sensitive to analysis approaches) seem to be necessary to interfere with the processing of returning echoes from echolocating bats (Kawahara and Barber (18)). In our data set, we find preliminary evidence of independent origins of sonar jamming in at least six moth subfamilies (Sphinginae, Macroglossinae, Aganainae, Arctiinae, Calpinae, Lymantriinae) based on this threshold. A seventh subfamily (Smerinthinae) also independently developed duty cycles capable of jamming, yet they are not capable of this behavior as this group lacks ears and thus cannot respond in advance to attacking bats. Animals that use complex tymbals with multiple microstriations (aganines, arctiines, and lymantrids) and stridulatory mechanisms (calpines and sphingids) are also likely capable of jamming. Thus, although moth morphology is not strictly deterministic of sound production function, some morphologies (wing beating mechanisms and tymbals with few microstriations; (44)) cannot support the high duty cycle (and likely high intensity) sounds necessary for jamming (18, 22).

Sonar jamming appears to be a derived strategy that has arisen repeatedly and recently in multiple lineages. Our preliminary investigations indicate that this strategy is not uniformly related to a loss (or lack of gain) of unpalatability to bats. We find that some genera capable of jamming bat sonar are palatable (Dataset S2; see Methods for palatability experimental details) and other genera are not, sometimes within the same subfamily (Arctiinae and Lymantriinae), thus the hypothesis that the origin of duty cycles capable of jamming frees lineages from the costs of sequestering chemicals for protection against bats (45) seems unlikely to be commonly supported. One possibility is that hostplant specialization canalizes sequestration strategies. Advertising difficulty of capture (evasive aposematism) is another conceivable function of conspicuous high duty cycle sounds (46) that may operate alongside sonar jamming, however, this hypothesis remains untested.

It appears that most sound-producing moths are not capable of jamming bat sonar. The majority of sound producers are therefore likely communicating with their bat predators, rather than disrupting echolocation. We found that moth genera that produced anti-bat sounds were commonly split between those that were palatable to bats and those that were not. Geometrid moths indeed seemed to be noxious, but not as repellent as lymantrids or arctiines (Dataset S2). Multiple subfamilies (Calpinae, Erebinae, Noctuinae, Nystaleinae, Macroglossinae, Smerinthinae, and Sphinginae) were considered quite palatable by the bats we pit these moths against (see supplement). These results likely indicate that these animals are exploiting the education imparted to their predators by unpalatable models (i.e., they are Batesian mimics).

To test the possible organization of anti-bat sounds into acoustic mimicry rings, we intensively studied a community of moths in Sumaco, Ecuador. We captured moths with UV lights and queried this megadiverse community for anti-bat acoustic response over 14 continuous nights. To analyze the resulting acoustic data, we used a dimensionality reduction algorithm (UMAP: Uniform Manifold Approximation and Projection; (47)) to find groups of moths with similar acoustic features (clusters). This unsupervised machine-learning algorithm estimates the topology of high dimensional data and uses this information to build a low dimensional representation that preserves relationships present in the data. We used 10 acoustic features (see Methods) and 33 species as input to UMAP to project the data from a 10-dimensional space into a 2D space where we found five well-separated clusters (Fig. 3; interactive 3D visualization at: http://projector.tensorflow.org/?config=https://raw.githubusercontent.com/nunezmatias/poli/main/ec6.json).

**Figure 3.**
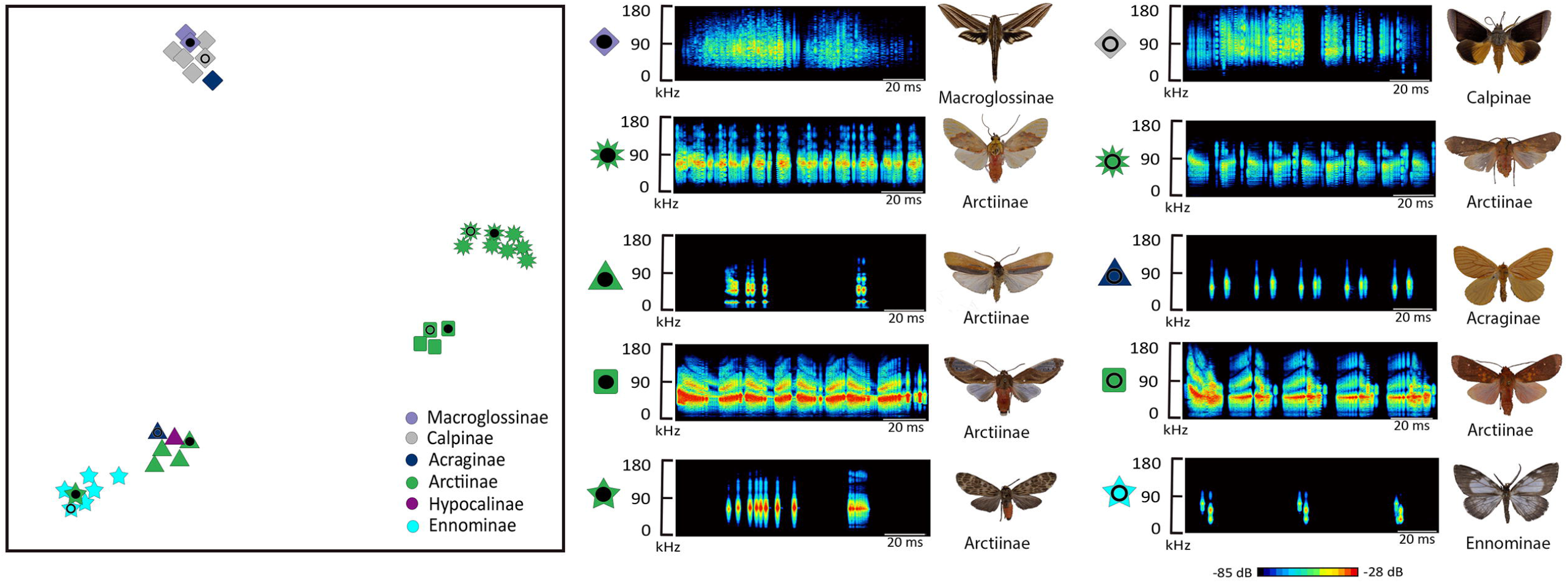
Purported acoustic mimicry rings of a community of moths in Sumaco, Ecuador (33 species). A UMAP (Uniform Manifold Approximation and Projection) projection shows clusters of moth anti-bat sounds with similar acoustic features. The relative distance between the clusters is meaningful in the sense that clusters that are close in the 2D map, are more similar than clusters that are further away. Photos of moths are congeners at the genus level. All photos taken by the authors. *Xylophanes titana*, purple diamond, solid circle; *Gonodonta syrna*, grey diamond, open circle; *Scaptius ditissima*, green sun, solid circle; *Melese sordida*, green sun, open circle; *Agylla* sp., green triangle, solid circle; *Acraga moorei*, dark-blue triangle, open circle; *Bertholdia bilineola*, green square, solid circle; *Melese chozeba*, green square, open circle; *Eucereon formosum dognini*, green star, solid circle; *Nephodia* sp., blue star, open circle. See Supplement Archive 11 for palatability data at the genus level.

While we caution that this analysis offers only a cursory temporal and spatial snapshot of the hyper-diverse mimetic associations that are likely present, we find some remarkable patterns. Each cluster of moth anti-bat sounds includes at least one species that we have found to be unpalatable to bats and most clusters also contain animals that bats readily consume. For example, one acoustic cluster contains one unpalatable dalcerid (Dalceridae), five palatable calpines (Erebidae: Calpinae), and two palatable sphingids (Sphingidae: Macroglossinae). Another cluster consists of six geometrid species (Ennominae) and one tiger moth (Erebidae: Arctiinae) all of which are likely honestly advertising noxious taste - perhaps a Müllerian ensemble. Interestingly, one cluster of Arctiini tiger moths (Erebidae: Arctiinae) uniformly contains extremely high duty cycle species capable of jamming bat sonar, including two genera that appear to be unpalatable to bats, supporting the prediction that jamming and aposematism are not mutually exclusive (27). Our preliminary data portends substantial community-level structuring of ultrasonic warning signals driven by the psychologies of syntopic bat predators (48). We are at the frontier of understanding a hidden dimension of biodiversity – the ultrasonic information transfer between bats and their insect prey.

Importantly, many species of moths also use ultrasonic sounds to transmit information to conspecifics – with males from at least six families (Crambidae, Erebidae, Geometridae, Noctuidae, Pyralidae, and Sphingidae) likely using this strategy to attract mates (49, 50). Some male moths use intense ultrasonic signals to communicate with females, as in tiger moths (Erebidae: Arctiinae) (50). Other families of moths produce quiet mating calls (Noctuidae, Arctiidae, Geometridae and Crambidae), apparently intended for nearby females (50). These “whispering” moths likely employ soft signals to avoid detection by eavesdropping bats and other predators (51–53). It is unclear if the use of ultrasound by moths evolved first in a mating context, or if it was secondarily co-opted from an anti-bat origin. Some moths are able to discriminate mates from bats, such as *Achroia grisella* (Pyralidae) females that exhibit differing behaviors, positive phonotaxis or freezing, when stimulated by different pulse rates (higher pulse rate indicating a conspecific calling male and lower pulse rate indicating an approaching bat, respectively (54). Alternatively, female *Spodoptera litura* (Noctuidae) are unable to distinguish attacking bats from ultrasound-producing males, suggesting a sensory exploitation origin of sound production in moths – that is, male moths exploit female freezing behavior to secure matings (55). We do not yet know whether moths that acoustically respond to echolocating bats are more likely to use ultrasound for mating, as many moths have not yet been tested for these behaviors (56), but this notion seems likely.

Ultrasonically-signaling moths appear to be connected by some of the most widespread and biodiverse mimicry complexes known to date (57, 58). The dynamics of these associations stand as a great unknown in natural history, and a laboratory for understanding mimicry dynamics and convergent evolution (59). The intense pressure to thwart the attacks of echolocating bats seems to have also driven ultrasound production in other insects. Tiger beetles (Cicindelidae) produce ultrasonic warning signals in response to sonar playback (60) and fireflies (Lampyridae: Lampyrinae), known to be noxious to bats (61), constantly produce ultrasonic clicks in flight, which may serve as a component of a multi-modal aposematic signal to bats (62). We predict that a complete understanding of ultrasonic mimicry rings will involve a thorough analysis of all major nocturnal, aerial insect groups including moths (Lepidoptera), beetles (Coleoptera), true bugs (Hemiptera), flies (Diptera), lacewings and antlions (Neuroptera) and more. Understanding how bat receivers generalize the massive numbers of insect warning sounds into categories is an important frontier in understanding this powerful selective force. Bats have shaped the nocturnal soundscape in profound ways – driving a chorus of nightly cries, across the globe, as moths and perhaps other insects jam sonar, warn of noxious chemicals, and mimic the sounds of unpalatable models. Comprehending this symphony is central to understanding insect biodiversity.

## Supporting information

.zip file of supplementary tables

## Acknowledgements

We thank the Danum Valley Conservation Area and Field Centre in Bornean Malaysia, Sabah Biodiversity Council, Dr. Chey Vun Khen of the Sabah Forestry Department; the Nouragues research field station in French Guiana (managed by CNRS) which benefits from “Investissement d’Avenir” grants managed by Agence Nationale de la Recherche (AnaEE France ANR-11-INBS-0001, Labex CEBA ANR-10-LABX-25-01) – with special thanks to Philippe Gaucher and Jerome Chave; Wildsumaco Biological Station in Ecuador – with special thanks to Jonas Nilsson; Gorongosa National Park and the E.O. Wilson Biodiversity Laboratory in Mozambique – with special thanks to Piotr Naskrecki, Marc Stalmans, James Byrne, Jason Denlinger, and Greg Carr. Kelly Dexter assisted with PCR. Kelly Dexter, Samantha Epstein and Kawahara lab volunteers prepared and organized voucher specimens. Ryan St Laurent for moth identification. Some analyses performed on the HiPerGator HPC (UF). We thank the National Science Foundation for funding (IOS-1920936 and IOS-1121807 to JRB; IOS 1920895 and IOS-1121739 to AYK). JJR supported in part by National Geographic Young Explorers Grant (YEG 9965-16). NTH supported in part by NSF GRFP award DGE-1315138. PRH was supported in part by National Geographic Society Waitt grant W318-14. The study was also funded by National Geographic Society grant CRE 9944-16 to JRB and two travel grants from the Nouragues research field station to JRB and AYK. We dedicate this work to the pioneering research of William Conner and James Fullard.

## Author Contributions

JRB and AYK designed and supervised the research and led all fieldwork. All authors collected data. DP led the phylogenetic analysis with input from AYK, JRB. MN led the machine learning analyses with input from JRB. JJR and KAM led moth sound analysis with input from JRB and assistance from BQ-R. DP and NTH led moth specimen identification. JRB and JJR wrote the first draft of the manuscript. All authors contributed to writing.

## Data and Materials Availability

The newly sequenced DNA barcodes used in this study have been deposited in the National Center for Biotechnology Information’s GenBank sequence database (all accession nos. provided in Dataset S4). All other data are available in the main text, the Supplementary Information, or at the Dryad Digital Repository (link to come when published).

## Methods

### Statement on Fieldwork Ethics

During our data collection trips, we received assistance, guidance, and hospitality from people in each of our field sites whose names we did not document. We recognize that this kind of expedition science is problematic and can be harmful to these communities in a variety of ways, including perpetuating colonial practices. In the future, we will strive to engage more deeply with the local population in the areas where we work and to offer more educational and professional opportunities. We remain indebted to those who helped us along this multi-year journey.

### Echolocation playback, tactile stimulation, and acoustic recording

We assayed moths in three of the world’s tropics: South America (Ecuador, French Guiana), Africa (Mozambique), and Asia (Malaysian Borneo) for ultrasonic reply to handling and bat attack. To simulate handling by a predator, we lightly compressed the moth’s head, abdomen, or thorax. We simulated bat attack using six recorded bat echolocation attack sequences (see supplement). Bat assemblages and echolocation strategies vary across the world. To capture some of the diversity of echolocation calls that moths might experience in different tropical regions, we presented moths with three different frequency modulated (FM) echolocation attacks and two constant frequency (CF) attacks. Two of the FM sequences were recorded from trained bats attacking a moth tethered 10 cm from a microphone (FM1: *Lasiurus borealis*, FM2: *Eptesicus fuscus*) (19). We also generated a synthetic bat attack based on the short-duration, broadband echolocation cries of some bats (63) (synthetic). To represent CF bat calls, we used on-board telemike recordings of bats (*Rhinolophus ferrumequinum nippon)* attacking prey provided to us by Yuki Kinoshita and Shizuko Hiryu (64) (CF1, CF2). All bat calls were played through an Avisoft UltraSoundGate Player BL Pro Speaker/ Amplifier (± 6 dB, 20-110 kHz, playback sampling rate 250 kHz) placed 10 cm behind the moth’s abdomen, except in the cases of sphingid moths, where the speaker was positioned on-axis 10 cm from the moth’s face, as their hearing organs are comprised of their mouthparts (65). Similarly, we recorded moth sounds using an Avisoft CM16 condenser microphone (±3 dB, 20-140 kHz) attached to an UltraSoundGate 116Hme DAQ sampling at 375 kHz via a laptop computer running Avisoft Recorder software, placed at a 90º angle 10cm from the moth’s thorax, except in the cases of sphingid moths, where the microphone was placed 10 cm directly behind the moth (as the genitals were previously known as the sound-producing organs in this group (19)).

Regardless of mechanism of ultrasound production, we focused our analyses on one complete modulation cycle of sound, which we defined as the two-component structure of the sound emissions. This paired structure results from: 1) the up-down wing stroke, 2) the buckling-unbuckling of tymbals, 3) the in-out or side-side stridulating of valves. We used Avisoft SASLab Pro software to measure three modulation cycles from each individual in our data set, except in cases where only two could be measured. We extracted the same parameters as those described in Barber & Conner (27) for comparability to other studies. To measure the temporal characteristics – duty cycle (proportion of 100ms window with moth sound present), duration of modulation cycle, and duration of modulation cycle components – we used the pulse train analysis tool with the following settings (Time constant=0.025ms, Threshold=0.15V, Hysteresis=15dB, Start/end threshold=-15dB, Envelope=Rectification + exponential decay, Pulse detection=Peak search with Hysteresis). We measured spectral characteristics – dominant frequency, frequency 15 dB above and below dominant frequency – from the Power Spectrum (averaged) tool with a Hann evaluation window and FFT=1024.

We attempted to record as many specimens as possible of each moth species, though this was usually limited by the number of healthy specimens we encountered in the field. For downstream analyses, we only considered a species to be responsive (i.e., producing ultrasound in response to bat ultrasound and/or tactile stimuli) if we recorded responsive ultrasound production in at least two specimens. Otherwise, the recorded species were assumed to be non-responsive. This is not the preferred method for obtaining negative data, since it is plausible that a moth could be capable of responding to stimuli, yet did not do so in our setting. However, we believed it was necessary to delineate between moths actually observed in the field, and moths that we were unable to test at all, but that were incorporated into our phylogeny. Thus, the non-responsive moths in the field were treated as having negative data, whereas the untested moths were treated as having missing data (see Phylogenetic Methods).

### Palatability

Palatability experiments were conducted on 93 moths from 26 species (see supplement) in the field. We ablated sound-producing structures (if present), before offering a hand-held captive bat (see supplement for species and locations) a moth via forceps. In an attempt to control for the foraging motivation of each bat, we only scored interactions where the bat was willing to eat a control moth (a species we knew to be palatable) both before and after we offered an experimental moth. We scored partial palatability by dividing the length of the moth body into six parts and assigning one point to the head, two points to the thorax, and three points to the abdomen, following the methods of Hristov and Conner (42). A palatability score of 0 indicates the moths was entirely rejected and a score of 6 indicates the moth was 100% consumed.

### Unsupervised machine learning cluster analysis of moth sounds

The dimensionality reduction algorithm Uniform Manifold Approximation and Projection (UMAP) (47) was used for finding groups of moth sounds with similar features (clusters). Dimensionality reduction algorithms capture variability in a limited number of random variables to allow two or three-dimensional visualization of data that resides in a multidimensional space. The most common approach is the method of principal component analysis (PCA) (66), which uses linear combinations of variables to generate orthogonal axes that capture the variation present in the data with fewer variables. Another approach, developed a century after PCA, t-Distributed Stochastic Neighbor Embedding (t-SNE) (67), carries out dimensionality reduction by analyzing similarity of points using a Gaussian distance in high dimensional space and mapping these data into a low dimensional space. t-SNE is able to capture local non-linear relationships in the data, which PCA by its linear design is not able to, but does not capture the global structure. A more recent method, UMAP, is an unsupervised machine-learning algorithm for dimension reduction based on manifold learning techniques and ideas from topological data analysis. It works by estimating the topology of the high dimensional data and uses this information to build a low dimensional representation that preserves relationships present in the data. It is better at mapping the global structure of the data from the high dimensional space than t-SNE, and is able to capture local relationships as well.

We used the moth acoustic features to define a multidimensional space where each moth is represented by a vector (or point) in that space. The data set consisted of 33 entries with 10 features each which translates to 33 points (vectors) in a 10-dimensional space. We input their coordinates into a PCA as a pre-processing step. The resultant principal components were then used as input to UMAP to project the data from the 10-dimensional space into a 2D space. Each cluster shares similar features. The relative distance between the clusters is meaningful in the sense that clusters that are close in the 2d map, are more “similar” that cluster that are farther away. The features variables used, extracted from audio files, were “MC DC mean”,“d MC mean”,“D 1/2 mean”,“D silent mean”,“D 2h mean”,“DF mean”,“D dB mean”,“+ 15 dB mean”,“-15 dB mean”,“100 ms DC mean” (see supplement for definitions). We used the software tools Scikit-learn (68) and pandas (69). The steps of dimensional reduction using the different methods we have discussed above can be seen in the interactive online version of the embedding (http://projector.tensorflow.org/?config=https://raw.githubusercontent.com/nunezmatias/poli/main/ec6.json) by clicking on the different bookmarks on the right (created via (70)).

### Phylogenetic methods

In order to determine the timing of evolution of anti-bat sound production in Lepidoptera, we created a dated molecular phylogeny, using the ages estimated in the Lepidoptera phylogeny of Kawahara et al. (12), that incorporates the moth taxa we tested for anti-bat ultrasound production. We attempted to find previously published COI barcodes and five commonly sequenced nuclear genes (CAD, DDC, EF1-A, period, wingless) for one species of every genus that was tested for anti-bat sound production (as well as the sound-producing genus tested in Corcoran and Hristov (20), and also used published data from as many species as possible that were included in the Kawahara et al. (12) dataset (this transcriptomic dataset lacked data for these six genes and thus could not directly be used). Whenever possible, molecular data for a genus was represented by a tested species; when such data were not available (after searching both NCBI and Bold Taxonomy Browser), a congener was used instead.

There were 11 genera from our sound production dataset that had no available sequence data; in order to represent these taxa in our analysis, we obtained new COI barcodes from DNA extracted from the legs of the ensonified specimens. DNA was extracted using an OmniPrep Genomic DNA Extraction Kit (G-Biosciences, St. Louis, MO), following the protocol of Espeland et al. (71) and PCR was performed following the protocol of Hebert et al. (72) using Lep1 reverse primers. Sanger sequencing was performed by Genewiz (South Plainfield, NJ). COI sequencing was unsuccessful for two non-sound-producing genera (*Grammodora, Trotonotus*), which were consequently excluded from the analysis. The nine newly sequenced barcodes used in this analysis were uploaded to NCBI ([GenBank IDs to be added after acceptance]), and specimen vouchers were deposited at the McGuire Center for Lepidoptera and Biodiversity (MGCL; Dataset S4). In total, our molecular dataset contained at least one gene for 432 Lepidoptera species.

Sequences for the six genes were aligned in MAFFT (73), then manually trimmed and concatenated in GENEIOUS v.11.1.5. The dataset was partitioned by codon position, constrained using the topology in figure 1 of Kawahara et al. (12), and a maximum likelihood analysis was performed in IQ-TREE v.1.6.2 (74), using ModelFinder to determine the best-fit substitution models for each partition (75). The resulting maximum likelihood tree was dated in TreePL (76), using the age estimates from Kawahara et al. (12) as secondary calibrations. The molecular dataset and other files associated with these analyses are included in Supplementary Archives 1–9.

Two ancestral state reconstructions (ASRs) of anti-bat sound production were performed using stochastic character mapping with the ‘make.simmap’ in the R package Phytools v07-70 (77). Symmetrical transition rate models were used in both ASRs, and 1000 simulations were performed. In order to reduce the amount of computational resources required, these ASRs were performed only on the Ditrysia clade of the dated tree, which comprise 93% of all taxa in the analysis (400/432). Only one non-Ditrysian genus had been tested for ultrasound production (Hepialidae: *Dalaca*, which did not produce ultrasound), so their absence did not significantly impact the ASR results since only 1/32 could have been confidently assigned a character state. In the first ASR, the evolution of anti-bat sound production was assessed by treating it as a ternary character, with taxa assigned to one of the following: 1. No sound production in response to a stimulus (this includes genera that constantly produce sound regardless of whether there is a stimulus, e.g. *Acraga*); 2. sound production in response to tactile stimuli; 3. sound production in response to both tactile stimuli and bat ultrasound (Dataset S5, Fig. S3). In instances where a species in the ensonified dataset was represented in the molecular dataset by a congener, we assumed that the congener had an identical character state. For taxa in the Kawahara et al. (12) dataset that were included in our ML analysis but not ensonified, an equal probability of 1/3 was assigned to each of the three states, if those taxa were known to have ears. For the untested taxa known to lack ears (12), we assumed they could not detect ultrasound and thus had no way to respond to bat calls, and we consequently assigned equal probabilities of 1/2 to the first two states, and 0 to the third state.

In the second ASR, the evolution of anti-bat sound production capable of jamming bat sonar (i.e., anti-bat ultrasound with a duty cycle value of at least 18%; (18)), was assessed by treating it as a binary character. Taxa were assigned to one of the following: 1. Duty cycle less than 18% (this includes genera that did not produce any ultrasound when tested); 2. Duty cycle of 18% or greater (Dataset S6, Fig. S4). As with the previous ASR, we assumed that congeners had identical character states. If duty cycle data were collected for multiple species in a genus, the value from the species with the largest mean duty cycle was used for that genus in the ASR (Supp. Archive 10). For untested taxa in the Kawahara et al. (12) dataset that were included in our ML analysis but not ensonified, an equal probability of 1/2 was assigned to each of the two states (regardless of whether they had ears). We also performed an ASR using maximum likelihood (‘anc.ML’ in Phytools v07-70 (77)), that modeled duty cycle as a continuous character (Dataset S7, Fig. S5). However, since this method cannot incorporate taxa with missing data, all non-ensonified taxa were assumed to have duty cycles of 0%.

