## Supplementary figures and legends for "Anti-Bat Ultrasound Production in Moths is Globally and Phylogenetically Widespread"

**Figure legends**

**Fig. S1.** Echolocation attack sequences used in playback experiments. To capture some of the diversity of echolocation calls that moths might experience in different tropical regions, we presented moths with three different frequency modulated (FM) echolocation attacks and two constant frequency (CF) attacks. Two of the FM sequences were recorded from trained bats attacking a moth tethered 10 cm from a microphone (FM1: *Lasiurus borealis,* FM2: *Eptesicus fuscus*). We also generated a synthetic bat attack based on the short-duration, broadband, frequency-modulated echolocation cries of some bats (synthetic). To represent CF bat calls, we used on-board telemike recordings of bats (*Rhinolophus ferrumequinum nippon)* attacking prey provided to us by Dr. Hiryu (CF1, CF2).

**Fig. S2.** Reproduction of Lepidoptera photos in Fig. 1 with numbers corresponding to column A in the photo accreditation table (Dataset S3).

**Fig. S3.** Ancestral state reconstruction of anti-bat ultrasound production in Lepidoptera. Black: No sound production in response to a stimulus; Orange: Sound production in response to tactile stimuli; Blue: sound production in response to both tactile stimuli and bat ultrasound.

**Fig. S4.** Ancestral state reconstruction of duty cycle associated with anti-bat sound production, treated as a binary character. Black: Duty cycle less than 18%; Red: Duty cycle at least 18%, implying ultrasound is capable of jamming bat sonar.

**Fig. S5.** Ancestral state reconstruction of duty cycle associated with anti-bat sound production, treated as a continuous character. Blue indicates the minimum duty cycle recorded in this study (0%) and red indicates the maximum (37%).

**Dataset legends**

**Dataset S1.** (provided as separate Excel file) List of Lepidoptera genera tested for anti-bat ultrasound production in this study. Asterisks indicate that ultrasound was observed, but was determined to not be in direct response to a stimulus. The genus *Eubaphe* (Geometridae: Larentiinae) could not be tested in this study and is absent from this list, but due to its known status as an anti-bat ultrasound producer (Corcoran and Hristov 2014), it is included in subsequent phylogenetic analyses and ancestral state reconstructions.

**Dataset S2.** (provided as separate Excel file) Subfamily averages of acoustic and palatability data from anti-bat responders. Mean values are followed by standard deviation in parentheses. *Total genera tested* – number of genera from each subfamily that were tested and were found to produce acoustic responses to bat echolocation sequences; *Mean MC DC* – mean modulation cycle duty cycle; *Mean MC Duration* – mean modulation cycle duration; *dB at dominant frequency* – amplitude of the most intense frequency in dB; *Mean Dominant Frequency* – mean frequency with the highest amplitude; *Bandwidth* range of frequencies 15 dB above and below dominant frequency; *Palatability* – palatability score ranging from 0 – 6 (uneaten – fully eaten; see Methods for greater detail) using the mean value of the averaged palatability score for each genus and only including genera that were found to respond to bat sonar; *Total genera offered* – number of genera in each subfamily that were offered to bats in palatability trials.

**Dataset S3.** (provided as separate Excel file) Sources of Lepidoptera photos used in Fig. 1. Numbers in column A correspond to the labeled photos in Fig. S2. All photos obtained from websites are licensed under Creative Commons licenses (see column H). Some subfamilies with ultrasound-producing genera are represented by a photograph of a non-sound producing genus; in these instances, the genus of the non-sound producer is indicated in column E.

**Dataset S4.** (provided as separate Excel file) GenBank IDs for specimens with newly sequenced barcode data. Vouchers for all specimens are deposited at the McGuire Center for Lepidoptera and Biodiversity (MGCL).

**Dataset S5.** (provided as separate Excel file) Character matrix for the ancestral state reconstruction used to study the evolution of anti-bat ultrasound production in Lepidoptera. Character states in each row are represented as probabilities summing to 1. For taxa included in the phylogenetic tree that were not ensonified, an equal probability of 1/3 was assigned to each of the three states, if those taxa were known to have ears. For non-ensonified taxa known to lack ears, a probability of 1/2 was assigned to the states in columns B and C.

**Dataset S6.** (provided as separate Excel file) Character matrix for the ancestral state reconstruction used to study the evolution of Lepidoptera ultrasound capable of jamming bat sonar (i.e., ultrasound with a mean duty cycle value of at least 18%). Duty cycle is scored as a binary character. For non-ensonified taxa, an equal probability of 1/2 was assigned to each of the two states (regardless of whether they had ears).

**Dataset S7.** (provided as separate Excel file) Character matrix for the ancestral state reconstruction used to study the evolution of Lepidoptera ultrasound capable of jamming bat sonar (i.e., ultrasound with a mean duty cycle value of at least 18%). Duty cycle is scored as a continuous character.

**Movie legends**

**Movie S1.** Ultrasound production by a male *Gonodonta bidens* (Erebidae: Calpinae). Movie shows the abdominal stridulatory mechanism from both a ventral view and a posterior view. Filmed at 120 fps and slowed down 4x.

**Movie S2.** Ultrasound production by a male *Lymantria* sp. (Erebidae: Lymantriinae) Movie shows a ventral view of the paired abdominal tymbals at the thoracic-abdominal junction. Filmed at 240 fps and slowed down 8x.

**Supplementary Archive 1. (provided as separate data file)**

Directory including .fasta files with raw Lepidoptera gene sequence data used to generate the molecular dataset for the phylogeny. All previously published sequences are labelled with their corresponding voucher numbers.

**Supplementary Archive 2. (provided as separate data file)**

Directory including .fasta files with aligned, trimmed, and renamed gene sequence data used to create the final concatenated molecular dataset for the phylogeny.

**Supplementary Archive 3. (provided as separate data file)**

Concatenated six-gene molecular dataset (in .fasta format).

**Supplementary Archive 4. (provided as separate data file)**

.tre file used to constrain the maximum likelihood (ML) phylogenetic analysis. Topology of the constraint tree is derived from the Lepidoptera phylogeny of Kawahara et al. (2019).

**Supplementary Archive 5. (provided as separate data file)**

Partitions and model selections (in .nex format) used when running the ML phylogenetic analysis in IQ-TREE.

**Supplementary Archive 6. (provided as separate data file)**

.tre file containing maximum likelihood best tree from the IQ-TREE analysis.

**Supplementary Archive 7. (provided as separate data file)**

Secondary calibrations (from Kawahara et al. 2019) used to date the ML phylogeny, formatted for input into a treePL .config file.

**Supplementary Archive 8. (provided as separate data file)**

.tre file with divergence time estimates from the treePL analysis, with ages of nodes listed in millions of years.

**Supplementary Archive 9. (provided as separate data file)**

Dated tree (.tre format) from Supplementary Archive 8 with non-Ditrysia species removed. This file was used for ancestral state reconstructions.

**Supplementary Archive 10. (provided as separate data file)**

Acoustic data for ultrasound-producing Lepidoptera species that were ensonified in this study.

**Supplementary Archive 11. (provided as separate data file)**

Genus-level palatability data for Lepidoptera fed to bats.

All Supplementary Archives are available at the Dryad Digital Repository (https://doi.org/??????)

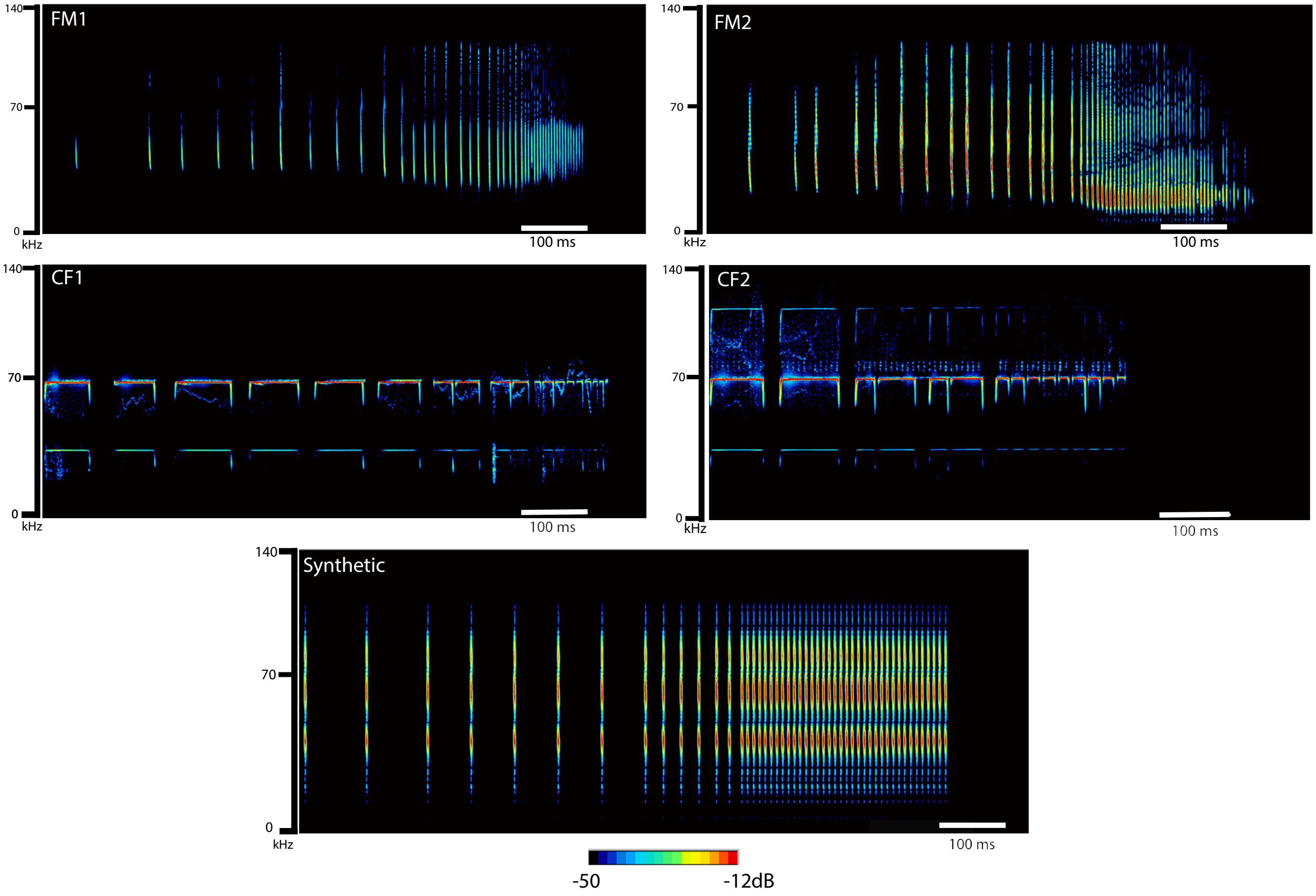


**Fig. S1.** Echolocation attack sequences used in playback experiments. To capture some of the diversity of echolocation calls that moths might experience in different tropical regions, we presented moths with three different frequency modulated (FM) echolocation attacks and two constant frequency (CF) attacks. Two of the FM sequences were recorded from trained bats attacking a moth tethered 10 cm from a microphone (FM1: *Lasiurus borealis*, FM2: *Eptesicus fuscus*). We also generated a synthetic bat attack based on the short-duration, broadband, frequency-modulated echolocation cries of some bats (synthetic). To represent CF bat calls, we used on-board telemike recordings of bats (*Rhinolophus ferrumequinum nippon*) attacking prey provided to us by Dr. Hiryu (CF1, CF2).

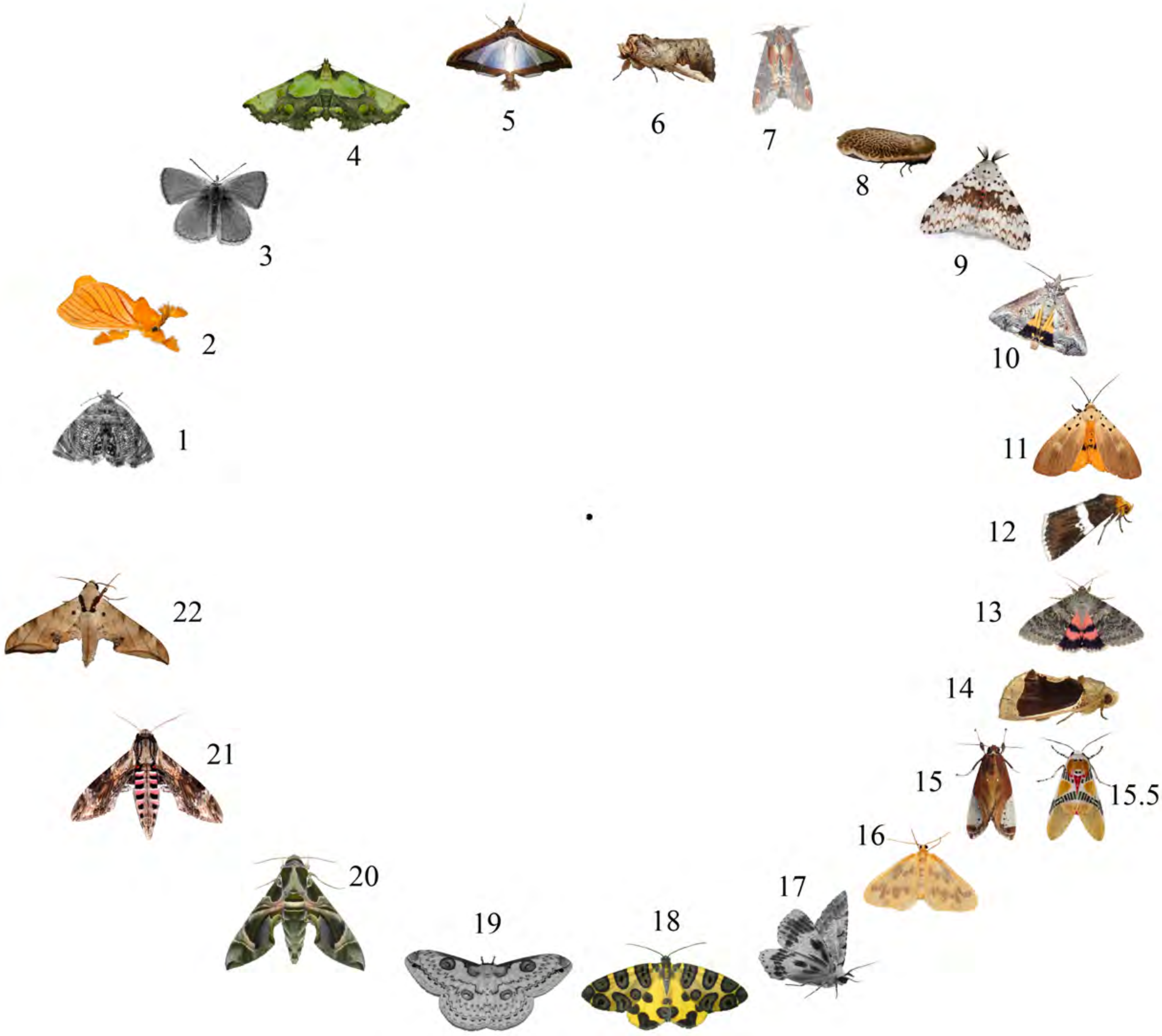


**Fig. S2.** Reproduction of Lepidoptera photos in Fig. 1 with numbers corresponding to column A in the photo accreditation table (Dataset S3).

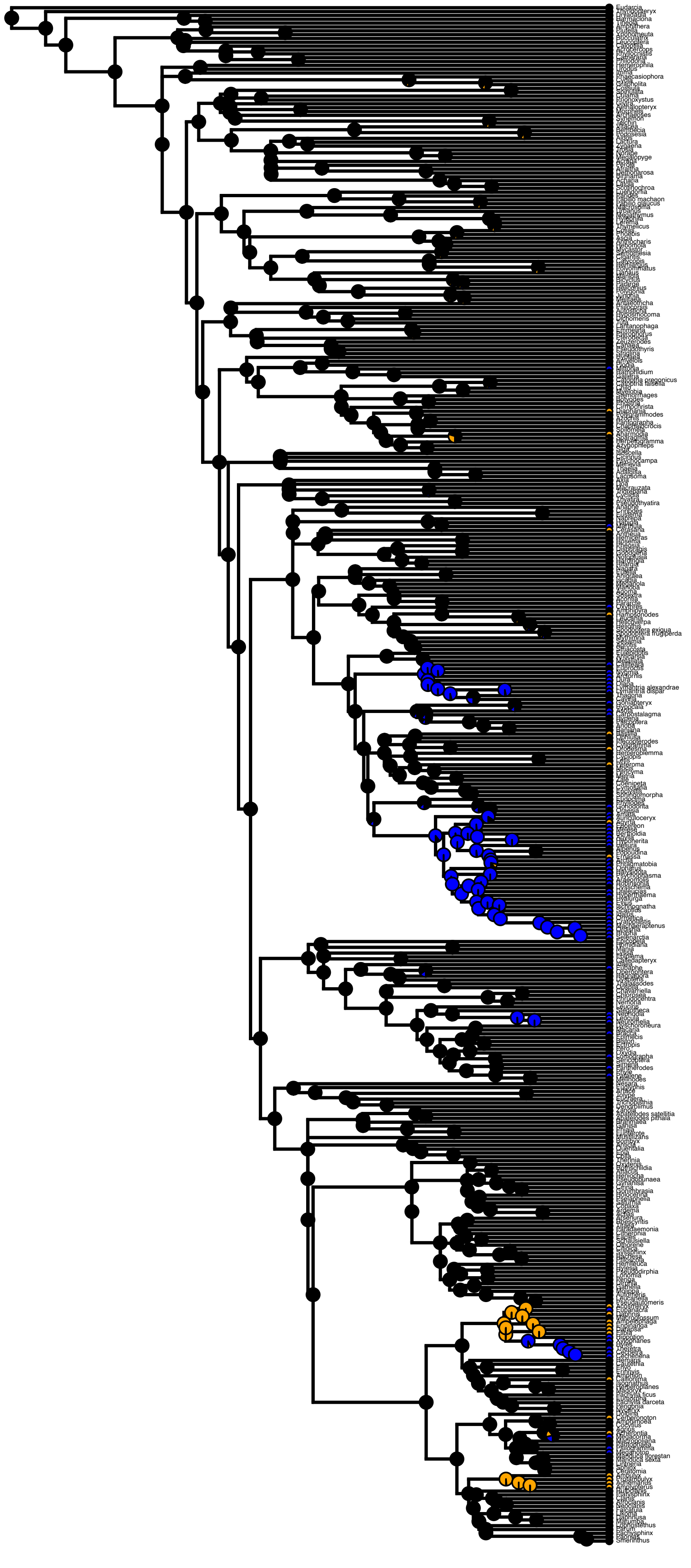

**Fig. S3.** Ancestral state reconstruction of anti-bat ultrasound production in Lepidoptera. Black: No sound production in response to a stimulus; Orange: Sound production in response to tactile stimuli; Blue: sound production in response to both tactile stimuli and bat ultrasound.

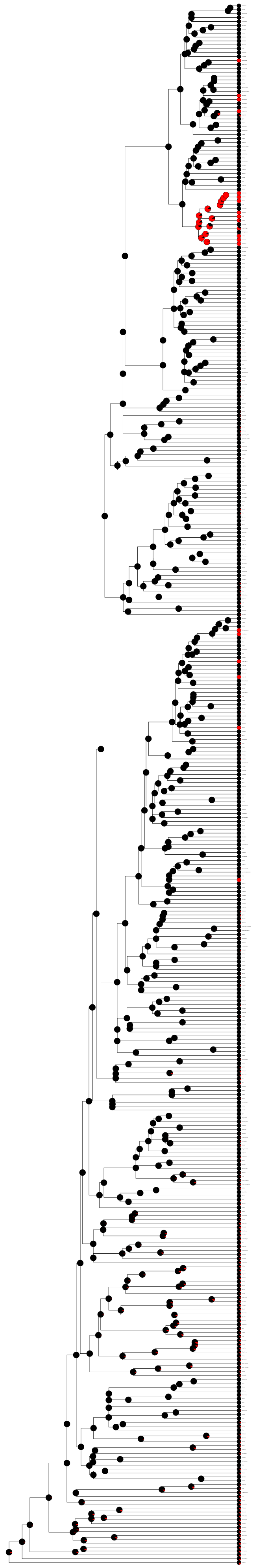

**Fig. S4.** Ancestral state reconstruction of duty cycle associated with anti-bat sound production, treated as a binary character. Black: Duty cycle less than 18%; Red: Duty cycle at least 18%, implying ultrasound is capable of jamming bat sonar.

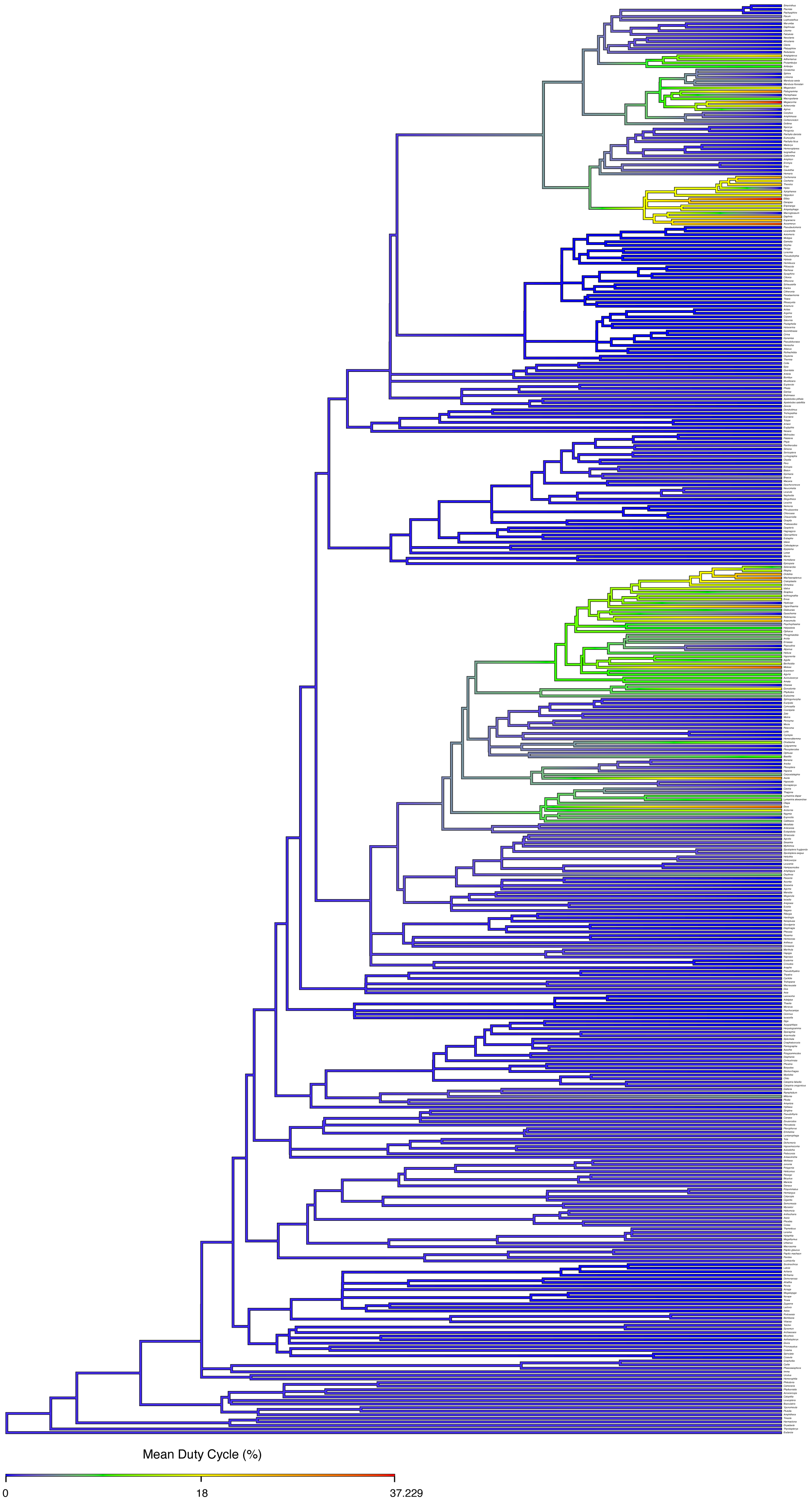


**Fig. S5.** Ancestral state reconstruction of duty cycle associated with anti-bat sound production, treated as a continuous character. Blue indicates the minimum duty cycle recorded in this study (0%) and red indicates the maximum (37%).
